# Probiotic Intervention Mitigates Radiation-Induced Intestinal Injury by Alleviating Oxidative Stress in a Human Gut-on-a-chip

**DOI:** 10.1101/2025.08.10.669539

**Authors:** Nam Than, Tadanori Mammoto, Donald E. Ingber, Hyun Jung Kim

## Abstract

Gastrointestinal acute radiation syndrome (GI-ARS) poses a critical public health concern, necessitating the development of effective medical countermeasures (MCM). Here, we evaluated the therapeutic potential of a commercially available probiotic formulation using human gut-on-a-chip model that mimics physiodynamic intestinal microenvironment. Intestinal epithelial Caco-2 cells were subjected to 8 Gray gamma radiation, targeting the epithelial layer, the culture medium, or both. The irradiated epithelial cells challenged to the irradiated medium resulted in significant DNA damage quantified by the presence of 53BP1 foci, increased cellular injury, and disrupted epithelial morphology. While epithelial irradiation alone did not compromise structural integrity, longitudinal exposure to irradiated medium induced oxidative stress, leading to morphological damage. Administration of the probiotic formulation significantly suppressed reactive oxygen species production, reduced epithelial damage, and preserved microarchitecture, independent of direct modulation of DNA damage. These findings suggest that probiotics may serve as promising live biotherapeutic MCM for mitigating GI-ARS in high-risk radiological exposures.

## INTRODUCTION

Exposure to ionizing radiation presents a serious health threat, particularly when high-energy radiation (e.g., gamma rays, X-rays, and neutrons) penetrates the body or when alpha and beta emitters are internalized through inhalation or ingestion^1,2^. One of the most severe outcomes of such exposure is acute radiation syndrome (ARS), occurring when the body receives a high dose of ionizing radiation above 1 Gray (Gy), such as during a nuclear accident, a radiotherapy overdose, or intentional total-body irradiation before bone marrow transplantation^3^. This level of exposure causes extensive damage to rapidly dividing cells, such as bone marrow and gastrointestinal (GI) cells^3^. Destruction of bone marrow stem cells leads to hematopoietic acute radiation syndrome (H-ARS), characterized by immunosuppression, and increased infection risk as a consequence^4^. Higher doses (> 6 Gy) can cause GI-ARS^5^, leading to severe diarrhea and fluid loss^6^, These symptoms are result from a loss of intestinal mucosa through oxidative stress and inflammation^7^, barrier dysfunction^7,8^, depletion of stem cells^9^, and immune impairment^10^. Although the U.S. Food and Drug Administration (FDA) has approved several therapies for H-ARS (e.g., filgrastim^11^, romiplostim^12^, and pegfilgrastim^13^), no treatments have yet been approved for GI-ARS because of a lack of human trials at this lethal dose and mechanistic understanding of drug target if performed on animals^14,15^. Therefore, the development of novel medical countermeasures (MCM) to protect or mitigate GI-ARS injury presents an unmet critical need.

Many radiomitigator drugs for GI-ARS in development aim at mitigating cellular damage and promoting cellular survival^14^. They have been demonstrated to inhibit apoptosis (e.g., Toll-like receptor 5 agonist CBLB502^16^), reduce inflammation (e.g., MIIST305^17^), promote crypt regeneration(e.g., glucagon-like peptide-2 analog [Gly2]GLP-2^18^), and neutralize reactive oxygen species (ROS) (e.g., CBLB502^19^). In this context, microbial therapies represent a strong addition to the MCM candidate portfolio for their recognized broad potential in reducing radiation-induced cell damage^20^. For example, *Lactobacillus*-containing probiotics could prevent side effects, such as diarrhea, during chemotherapy^21^. *Lacticaseibacillus rhamnosus* GG (LGG) was found to released lipoteichoic acid that agonized Toll-like receptor (TLR)-2, which stimulated cyclooxygenase-2 (COX-2)-positive mesenchymal stem cells to migrate to injured regions and promote crypt survival and reduce apoptosis in mice following a 12 Gy whole body irradiation^22,23^. These supporting evidence led to an ongoing FDA-approved phase-3 clinical study (NCT01790035) to study the effects of LGG in GI cancer patients undergoing chemoradiation. Commercially available formulations like VSL#3, which contain 8 beneficial bacterial strains, have also demonstrated the ability to mitigate radiation-induced diarrhea in GI cancer patients undergoing radiation therapy, although its mechanism was not elucidated^24^. Studies of chemotherapy-induced diarrhea in mice have attributed the radioprotective effects of VSL#3 to crypt proliferation and inhibition of apoptosis^25^. However, the ROS-neutralizing effect of VSL#3 has been underexplored in GI-ARS, even though its anti-inflammatory and antioxidant effects have been demonstrated on various non-radiation diseases^26–28^.

Traditional MCM research relies heavily on animal models to establish dose-response relationships between radiation and tissue damage, crypt loss, and survival outcomes^29^. These models were used to evaluate the effectiveness of various treatments such as hematopoietic growth factors^30^, antioxidants^31^, radioprotectors^32^, and probiotics^33^. However, there is consensus that translating findings from animal models to humans remains difficult due to significant physiological differences, such as faster epithelial turnover rate in mice^34^, distinct microbial composition^35^, insusceptibility of mice to human infection^36^, and varying cytokine profiles^37^. While non-human primate models provide physiological similarity and key features of human GI-ARS response, such as rapid stem cell loss, villus atrophy, ulceration, and barrier dysfunction^38^, their use is ethically contentious and financially restrictive^39^. More importantly, ROS are key mediators of radiation-induced damage^40^, but measuring such transient molecules *in vivo* is challenging^41^.

Conventional *in vitro* models, such as Transwells and organoids, are limited by short co-culture durations, bacterial overgrowth, and a lack of oxygen control^42–44^. Organoids require microinjection for microbial access, restricting long-term studies^45^. To overcome these limitations, the human gut-on-a-chip was developed to replicate key features of the intestinal microenvironment, including luminal flow, peristalsis-like motion, and oxygen gradients^46–49^. Using this model, we previously identified radiation-induced epithelial injury and demonstrated the therapeutic potential of a prolyl hydroxylase inhibitor (DMOG)^50^. The gut-on-a-chip now provides a robust platform to evaluate probiotics as candidate MCMs for GI-ARS. Importantly, this dynamic microenvironment can recreate physiologically relevant conditions that support stable microbial colonization without overgrowth, while maintaining chemical homeostasis and steady-state nutrient supply^46,51,52^. This advancement is particularly crucial for evaluating the therapeutic potential of human probiotics as a GI-ARS MCM, specifically aimed at mitigating oxidative stress.

In this study, we investigated the protective effects of a human-derived probiotic formulation (VSL#3) against radiation-induced intestinal injury using a physiologically relevant human gut-on-a-chip model. By independently irradiating cellular and acellular components (e.g., culture medium), we delineated their distinct contributions to nuclear damage and intracellular oxidative stress. The probiotic treatment significantly reduced intracellular ROS levels and preserved epithelial integrity under radiation challenge. These findings highlight the potential of live biotherapeutic agents as promising candidates of MCM for GI ARS.

## RESULTS

### Gamma irradiation induces intestinal epithelial injury and DNA damage

We utilized a gut-on-a-chip microfluidic platform constructed from optically transparent and elastic polydimethylsiloxane (PDMS), comprising two parallel microchannels separated by a porous PDMS membrane to establish distinct luminal and abluminal compartments^46^. Human intestinal epithelial Caco-2 cells seeded onto the extracellular matrix (ECM)-coated luminal surface of the membrane and cultured under continuous perfusion (30 µL/h) for approximately 100 h, promoting the development of a 3D epithelial^48,49^ (Figure 1A, left schematic). To mimic intestinal peristalsis, the device was subjected to cyclic mechanical strain (10% deformation at 0.15 Hz frequency), replicating physiodynamic mechanical cues^51^. To model GI-ARS *in vitro*, we exposed the entire chip system, including both 3D epithelial cell layers and culture medium, to high-dose gamma irradiation^53^ (total 8 Gy at 0.78 Gy/min) (Figure S1A), followed by perfusion of irradiated medium under continued mechanical actuation for an additional 72 h (Figure 1A). This treatment resulted in marked epithelial injury, including significant disruption of the 3D epithelial architecture and loss of cellular contours (Figures 1B and S1B). To quantify cellular damage, we measured extracellular lactate dehydrogenase (LDH) activity. The irradiated group exhibited a significant elevation of LDH release (70.4±8.62%) in the apical microchannel, representing an approximate 3.8-fold increase compared to non-irradiated controls (*p*<0.01) (Figure 1C, AP). In contrast, LDH release from the basolateral compartment remained minimal (Figure 1C, BL), suggesting polarized epithelial injury. These results demonstrate that intestinal epithelial cells within the PDMS-based gut-on-a-chip system are highly susceptible to high-dose radiation, recapitulating key features of GI-ARS.

**Figure 1.**
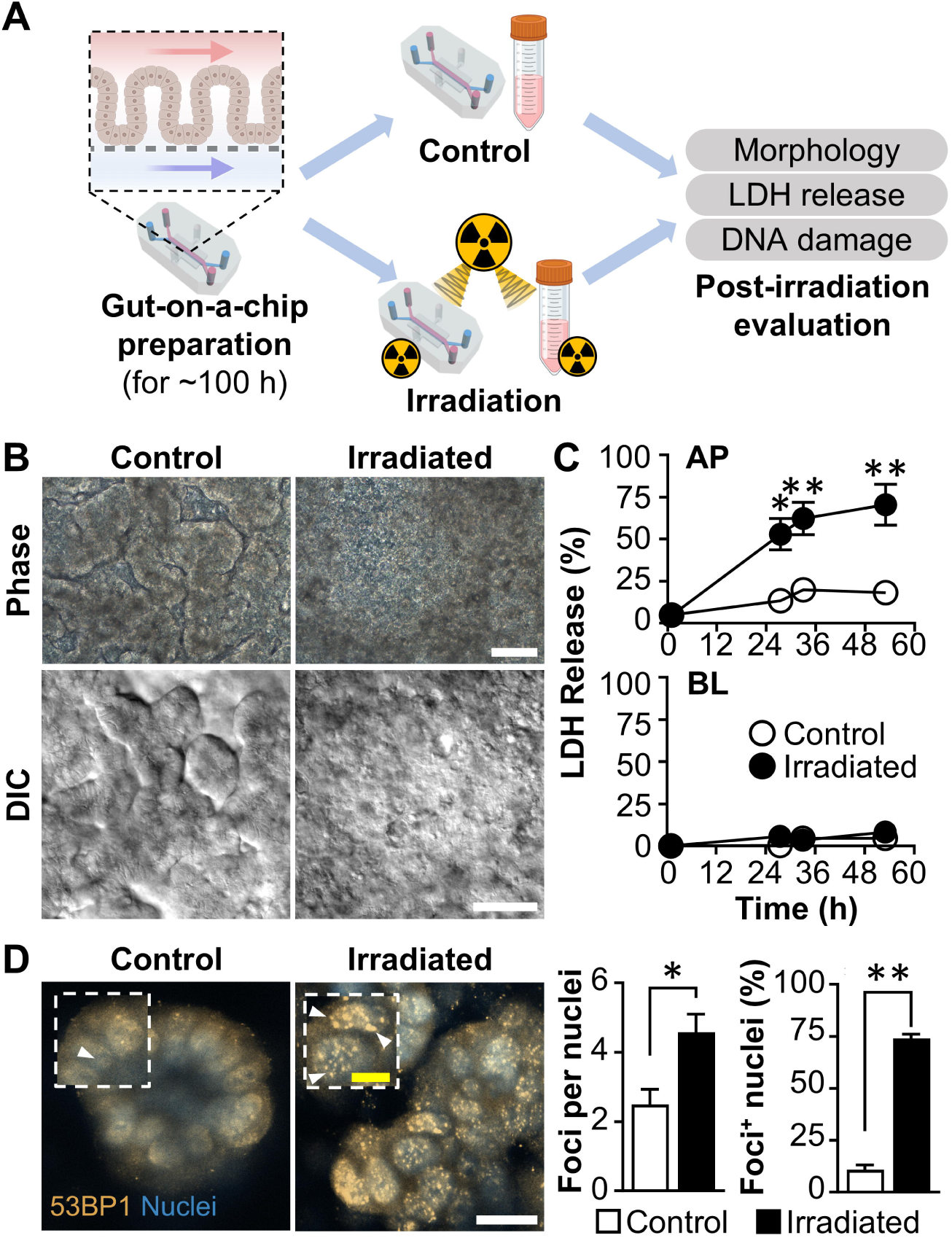
Gamma irradiation-induced epithelial injury and DNA damage in a gut-on-a-chip. (A) Schematic representation and experimental timeline illustrating the gut-on-a-chip setup used for gamma irradiation studies. Irradiation icons indicate the targeted components, either the epithelial cells within the chip or the culture medium, subjected to 8 Gy exposure, followed by subsequent cellular and biochemical analyses. (B) Representative micrographs showing the morphology of Caco-2 intestinal epithelial cells cultured in a gut-on-a-chip under control (non-irradiated) or irradiated (8 Gy) conditions. Cells were continuously perfused with irradiated medium (8 Gy; 30 µL/h) and subjected to cyclic mechanical strain (10% deformation, 0.15 Hz frequency) for 72 h. Images were acquired using phase-contrast (top) and differential interference contrast (DIC; bottom) microscopy. Bars, 100 μm. (C) Quantitative analysis of cytotoxicity based on lactate dehydrogenase (LDH) release in apical (AP) and basolateral (BL) compartments under irradiated conditions (8 Gy) with irradiated medium perfusion (30 µL/h) and mechanical strain (10%, 0.15 Hz) in a gut-on-a-chip. Data represent mean ± SEM (*n*=2). *\*p*<0.05; *\*\*p*<0.01. (D) Immunofluorescent detection of DNA double-strand breaks via 53BP1 nuclear foci (arrowheads) in control and irradiated gut-on-a-chip systems, 5 hours post-radiation exposure. Confocal micrographs are accompanied by quantitative analysis of the average number of 53BP1 foci per nucleus and the percentage of foci-positive nuclei (*n*=7-14). Yellow bar, 10 μm; White bar, 20 μm. *\*p*<0.05; *\*\*p*<0.0001.

We next conducted a spatial analysis of tumor protein p53-binding protein 1 (53BP1), a marker of DNA double-strand breaks (DSBs), to evaluate nuclear damage following gamma radiation exposure^54^. Immunofluorescence microscopy revealed punctate nuclear foci of 53BP1 in Caco-2 cell monolayers subjected to irradiation, whereas non-irradiated controls exhibited diffuse nuclear 53BP1 distribution (Figure S2). Similar nuclear foci patterns were observed in our microfluidic system as early as 5 h post-exposure to 8 Gy of gamma radiation (Figure 1D, arrowheads). Quantitative analysis showed a significant increase in both the number of 53BP1 foci per nucleus (1.8-fold, *p*<0.05) and the percentage of foci-positive nuclei (6.8-fold, *p*<0.0001) in irradiated conditions compared to non-irradiated controls. These findings indicate that exposure to radiation directly to epithelial cells followed by the perfusion of irradiated culture medium induces pronounced DNA damage, morphological disruption, and nuclear relocalization of 53BP1, confirming the cellular injury response in our model.

### Irradiated medium alone can cause epithelial damage and morphological disruption

To delineate whether the observed epithelial injury was attributable solely to direct irradiation of the cells or also involved indirect effects mediated by the surrounding microenvironment, we leveraged the modular design of the gut-on-a-chip platform to independently assess the contributions of irradiated components. Three experimental conditions were established: i) simultaneous irradiation of both the epithelial cells within the device and the culture medium (“Irradiated All”), ii) irradiation of epithelial cells followed by perfusion with fresh, non-irradiated medium (“Irradiated Cells”), and iii) perfusion of irradiated medium into a device containing non-irradiated epithelial cells (“Irradiated Medium”) (Figure 2A). Morphological assessment revealed substantial disruption of the epithelial microarchitecture in both the “Irradiated All” and “Irradiated Medium” groups, whereas the “Irradiated Cells” group retained intact 3D epithelial structure when conditioned to non-irradiated fresh medium. These findings suggest that irradiation of the culture medium alone is both necessary and sufficient to induce substantial 3D morphological deterioration of the intestinal epithelium within the chip.

**Figure 2.**
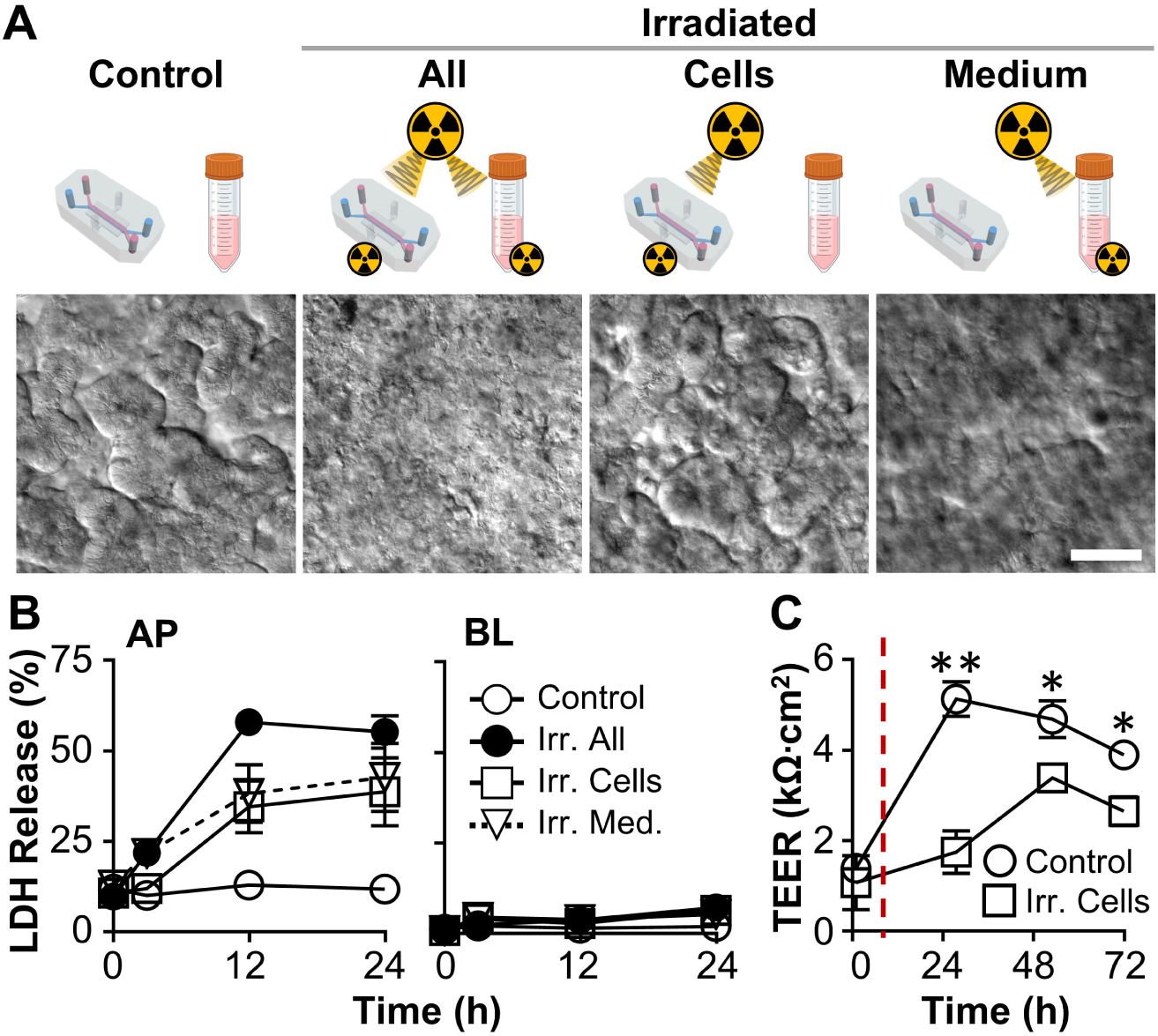
Irradiated culture medium alone induced epithelial injury and barrier dysfunction in a gut-on-a-chip. (A) Schematic overview of experimental conditions and corresponding epithelial morphologies observed in gut-on-a-chip cultures 72 hours post-exposure. Groups included non-irradiated control, full irradiation of both cells and medium (Irr. All), irradiation of epithelial cells alone (Irr. Cells), and irradiation of medium alone (Irr. Medium). Representative DIC images show morphological alterations across conditions. Bar, 100 μm. (B) Quantification of cytotoxicity assessed by LDH release in the apical (AP) and basolateral (BL) compartments under each experimental condition. All chips were cultured under continuous flow (30 µL/h) and cyclic mechanical strain (10% cell strain, 0.15 Hz). Data are represented as mean ± SEM (*n*=3-12). All data points for the “Irr. All,” “Irr. Cells,” and “Irr. Medium” groups in the AP compartment at both 24 and 48 h demonstrated statistically significant differences compared to the non-irradiated control with *p*<0.01. However, no significant differences were observed between the “Irr. Cells” and “Irr. Medium” groups at these time points. (C) Assessment of epithelial barrier function over time using TEER measurements following exposure to irradiated medium for 90 h (*n*=4). The red dashed line indicates the time point of irradiation. *\*p*<0.05; *\*\*p*<0.001.

To evaluate the impact of distinct irradiation strategies on epithelial cytotoxicity, we quantified extracellular LDH levels from both AP and BL compartments as a function of time. All irradiated conditions exhibited significantly elevated LDH release in the AP channel compared to non-irradiated controls, displaying the highest LDH levels throughout the observation period, indicative of the most severe cytotoxic response (Figure 2B). Both the “Irradiated Medium” and “Irradiated Cells” groups showed comparable increases in LDH release, reaching approximately 40% by 48 h without statistical significance, suggesting that the extent of cellular injury is not strictly aligned with observed morphological disruption.

Consistent with prior observations, LDH release remained negligible in the BL compartment. Given the notable cytotoxicity induced by irradiated medium alone, we next assessed epithelial barrier integrity using transepithelial electrical resistance (TEER) as a functional readout. In control chips, TEER values rapidly increased to >5 kΩ·cm^2^ within 30 h, reflecting the establishment of intact tight junctions, followed by a gradual decline to ∼4 kΩ·cm^2^ up to 72 h. In contrast, epithelial cells exposed to irradiated medium exhibited a markedly delayed TEER elevation, reaching only 3.38 kΩ·cm^2^ at 50 h, significantly lower than controls (*p*<0.05; Figure 2C). These results demonstrate that irradiated medium impairs both epithelial viability and functional barrier restoration in a gut-on-a-chip model.

### Irradiated medium can induce epithelial ROS without DNA damage

Given the pronounced impact of irradiated medium on epithelial morphology and viability, we further examined its role in inducing ROS generation and DNA damage, key cellular responses associated with radiation exposure^40^. Non-irradiated epithelial cells perfused with irradiated medium under dynamic flow and mechanical strain for 72 h exhibited markedly elevated intracellular ROS levels, as measured by CellROX fluorescence (5 μM), demonstrating approximately a 15-fold increase compared to non-irradiated controls (*p*<0.0001; Figure 3A). In parallel, extracellular ROS levels quantified from the culture medium using H2DCFDA (5 μM) showed a ∼4-fold increase after just 1 h of exposure to irradiated medium (*p*<0.0001; Figure 3B), indicating a rapid oxidative response. Despite the heightened oxidative stress, epithelial cells exposed to irradiated medium for 72 h did not exhibit a significant increase in the number of 53BP1 nuclear foci per cell (Figure 3C, left panel). However, a modest but significant elevation in the proportion of foci-positive nuclei was observed relative to the control group (Figure 3C, right panel; *p*<0.01). Noticeably, the foci-positive cells in the “Irr. Medium” group were on the outer edge of the 3D microstructure where cells were in direct contact with the ROS-rich irradiated medium. The results suggest that while irradiated medium induces considerable oxidative stress in most cells, its capacity to elicit overt DNA double-strand breaks is correlated to the spatial distance of contact.

**Figure 3.**
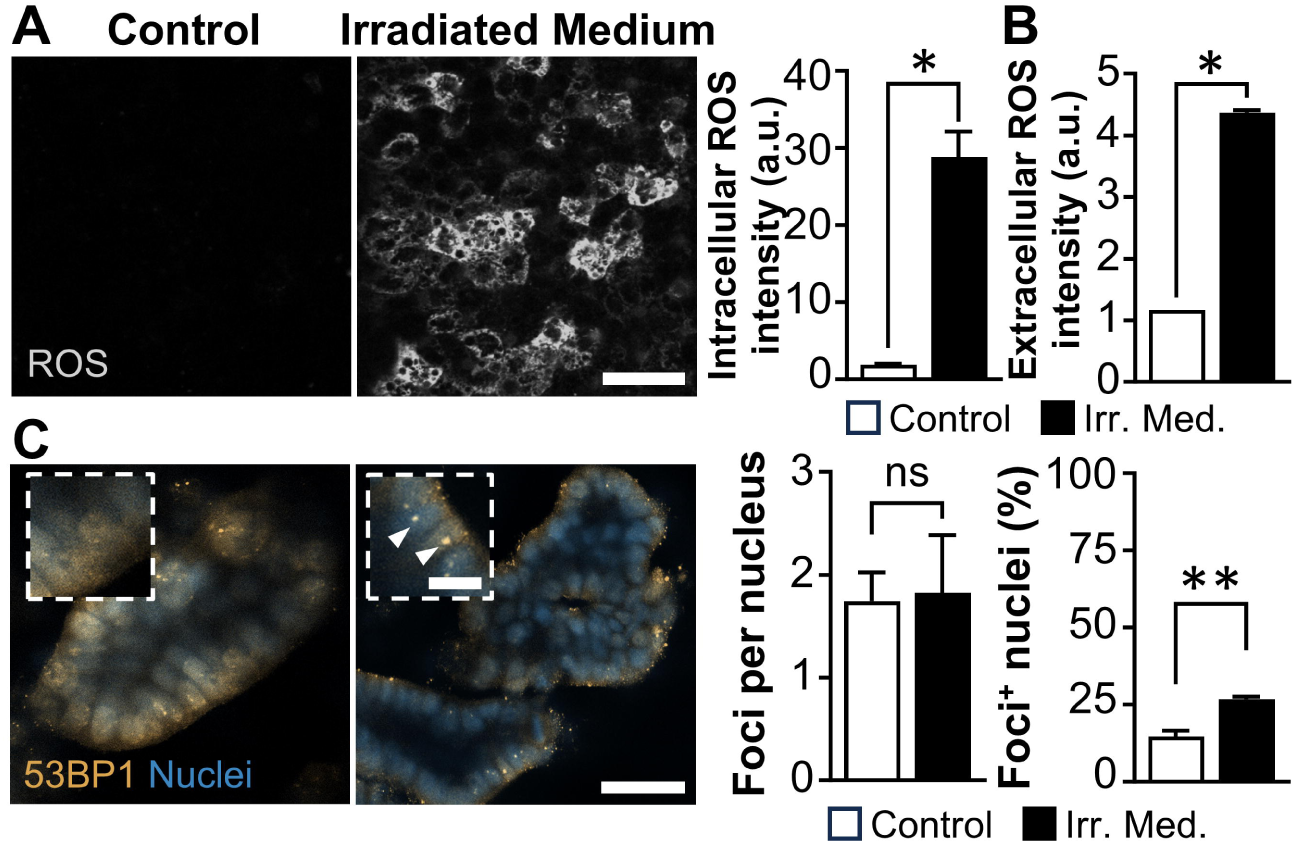
Irradiated medium induces oxidative stress independent of overt DNA damage. (A) Assessment of intracellular ROS levels in epithelial cells following 72-h exposure to irradiated medium. ROS were visualized using fluorescent ROS probes and imaged via confocal microscopy. Representative micrographs were provided alongside quantification of fluorescence intensity (*n*=10). Bar, 50 μm. *\*p*<0.0001. (B) Quantification of extracellular ROS production using the H2DCFDA assay in epithelial cell cultures challenged with the irradiated medium for 72 h. Data are presented as mean ± SEM (*n*=3). *\*p*<0.0001. (C) Evaluation of nuclear DNA damage via immunofluorescent detection of 53BP1 foci, indicative of double-strand DNA breaks, in epithelial cells exposed to irradiated medium for 5 h. Confocal images highlight foci localization (arrowheads), accompanied by quantification of average foci per nucleus and percentage of foci-positive nuclei (*n*=4-10). *\*p*<0.01; ns, not significant. Yellow bar, 20 μm; White bars, 50 μm.

### Probiotic bacteria ameliorate radiation-induced injury

To evaluate the therapeutic potential of probiotics against radiation-induced intestinal injury, we administered a live probiotic formulation, VSL#3, enriched *in vitro* (final density, 1×10^7^ CFU/mL; MOI, 2.5) into the luminal channel of the gut-on-a-chip device either prior to irradiation (Pre-VSL#3) or post-irradiation (Post-VSL#3) (Figure 4A). As previously observed, epithelial cells exposed to irradiated medium in the absence of microbial cells (Germ-free control) exhibited severe morphological deterioration, including a complete loss of epithelial structure within 72 h (Figure 4B). In contrast, epithelial co-culture with VSL#3 prior to gamma irradiation markedly preserved epithelial 3D microstructure, particularly in chips where non-irradiated epithelial cells were challenged with irradiated medium. In these cases, the epithelial contours and apical protrusions remained visibly intact, indicating a robust protective effect.

**Figure 4.**
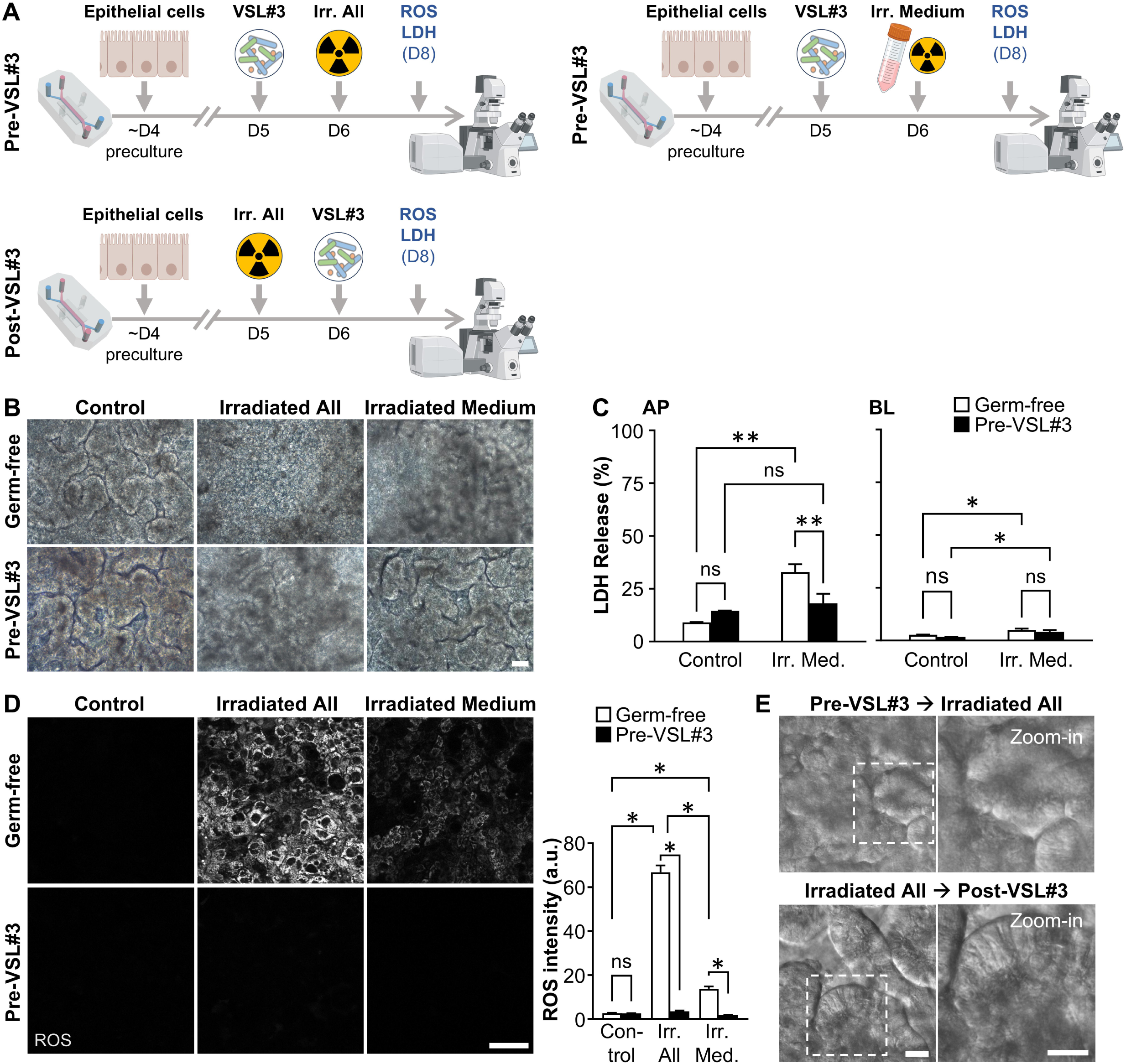
Probiotic VSL#3 attenuated radiation-induced epithelial injury in a gut-on-a-chip. (A) Schematic overview of the experimental workflow. “Pre-VSL#3” and “Post-VSL#3” denote the timing of probiotic administration, either prior to or following gamma irradiation of both epithelial cells and culture medium. Co-culture with VSL#3 was initiated once Caco-2 epithelial cells established a mature 3D architecture under flow (40 µL/h) and mechanical strain (10% deformation, 0.15 Hz). Post-exposure assessments included intracellular ROS quantification via microfluorimetry and cytotoxicity evaluation through LDH assays conducted on culture supernatants. (B) Phase-contrast micrographs showing epithelial morphology in non-irradiated controls, “Irr. All,” and “Irr. Medium” conditions, with or without VSL#3 pre-treatment. VSL#3 was pre-administered for 24 h before irradiation treatments and continued to be co-cultured with epithelial cells for 72 h post-irradiation. Bar, 50 μm. (C) Quantification of cytotoxicity, assessed by LDH release in AP and BL compartments across experimental conditions, including non-irradiated and germ-free controls, with or without VSL#3 pre-treatment. All chips were operated under continuous flow (40 µL/h) and cyclic mechanical deformations (10%, 0.15 Hz). Data are represented as mean ± SEM (*n*=2-4). *\*p*<0.01; *\*\*p*<0.001; *\*\*\*p*<0.0001; ns, not significant. (D) Evaluation of intracellular ROS accumulation in epithelial cells, irradiated or non-irradiated, following 72-h exposure to irradiated medium, with or without VSL#3 co-cultures. Confocal fluorescence micrographs and quantification of signal intensity are shown (*n*=2–10). Bar, 50 μm. *\*p*<0.0001. (E) DIC images depicting epithelial morphology 48 h post-irradiation and irradiated medium perfusion, with either pre-or post-treatment with VSL#3. Pre-treatment preserved epithelial contours, while post-treatment further enhanced structural recovery. Bars, 25 μm.

Quantification of cytotoxicity using LDH assays revealed that VSL#3 treatment significantly reduced LDH release into the apical channel when cells were challenged with irradiated medium, demonstrating a ∼1.8-fold reduction compared to the germ-free irradiated control (*p*<0.001; Figure 4C, AP). A slight, though statistically insignificant, increase in LDH levels was noted in the VSL#3-only control group (Figure 4C, AP), potentially reflecting a baseline LDH release by probiotic microbes under irradiation (Figure S3). Basolateral LDH levels remained negligible across all groups (Figure 4C, BL).

Consistent with previous findings, the irradiated medium induced robust intracellular ROS production regardless of whether epithelial cells were directly irradiated, under germ-free conditions (Figure 4D). However, co-culture with VSL#3 significantly suppressed ROS generation in both the “Irr. All” (19.3-fold reduction, *p*<0.0001) and “Irr. Medium” (7.3-fold reduction, *p*<0.0001) groups compared to their respective germ-free irradiated controls (Figure 4D, Pre-VSL#3). We confirmed that cultured VSL#3 cells exhibited significantly greater extracellular ROS scavenging functions than the germ-free control across hydrogen peroxide concentrations up to 1 µM (Figure S4A), indicating that the observed reduction in ROS in both Irr. All and Irr. Medium conditions was likely attributable to the ROS-mitigating activity of VSL#3.

In addition, VSL#3 cells readily exhibited a peroxidase activity of approximately 2 mU/mL within 30 min when challenged with 2 mM H_2_O_2_ (Figure S4B). Interestingly, administration of VSL#3 after irradiation (Post-VSL#3) resulted in improved preservation of epithelial morphology, with columnar epithelial structures clearly maintained (Figure 4E), suggesting that VSL#3 may exert both protective and reparative effects against radiation-induced injury.

Despite these protective effects on viability and oxidative stress, VSL#3 did not mitigate DNA damage in the “Irr. All” group, as evidenced by similar levels of 53BP1 foci per nucleus (2.90±0.31 vs. 2.78±0.26; *p*>0.05) and comparable percentages of foci-positive nuclei (97.53±0.41% vs. 94.93±0.92%; *p*>0.05) relative to the germ-free irradiated control (Figures 5 & S5). However, in the “Irr. Medium” condition, VSL#3 pre-treatment significantly reduced the average number of 53BP1 foci per nucleus (4.2-fold reduction; *p*<0.01) and the proportion of foci-positive nuclei (2.7-fold reduction; *p*<0.0001), indicating partial mitigation of DNA damage when direct irradiation was absent. Similar to prior observations, the foci-positive cells were on the outer side of the 3D microstructure of the “Irr. Medium” group.

**Figure 5.**
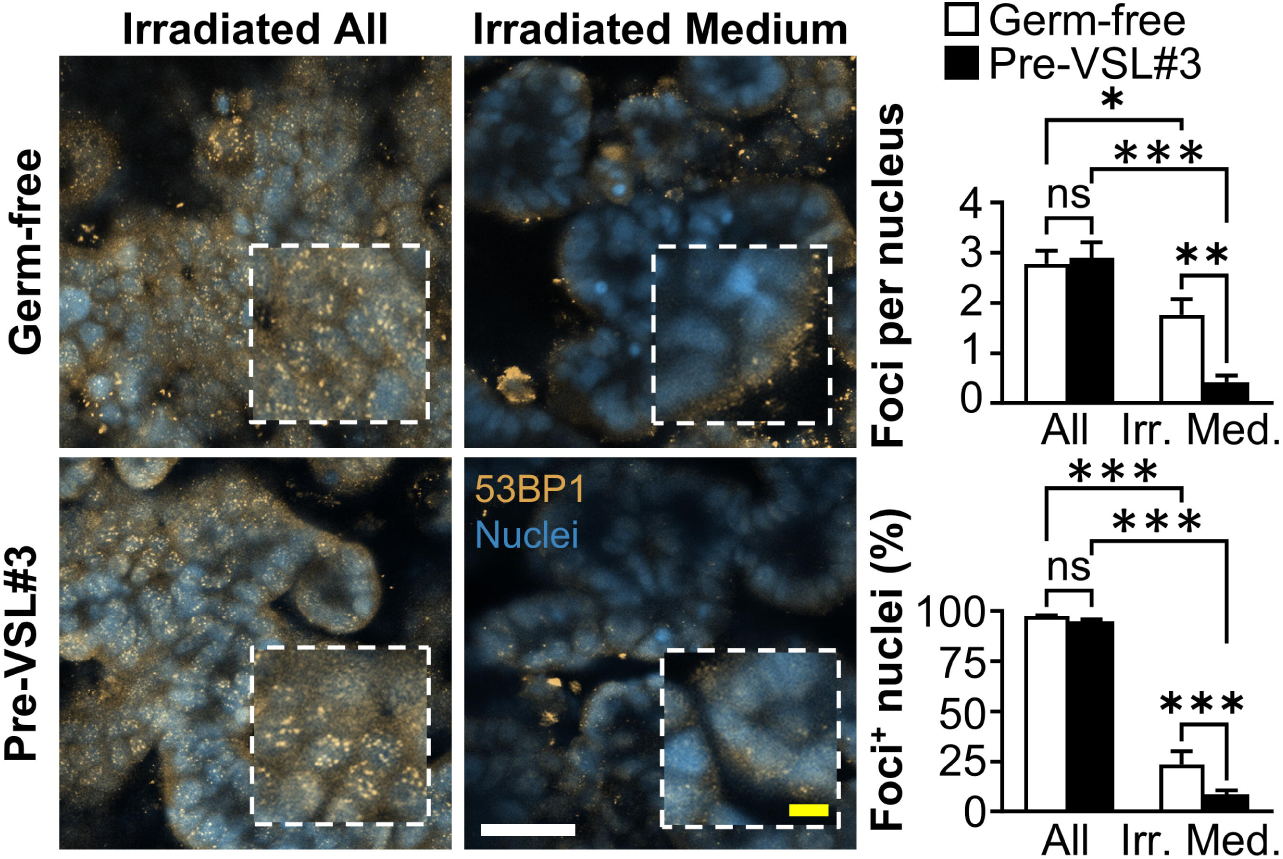
Probiotic VSL#3 did not modulate radiation-induced DNA damage response. Immunofluorescent detection of nuclear double-strand DNA breaks was performed using 53BP1 staining in gut-on-a-chip cultures subjected to full irradiation (“Irr. All”) or irradiated medium only (“Irr. Medium”), with or without VSL#3 pre-treatment. Confocal micrographs were captured 5 h post-irradiation under continuous flow (40 µL/h) and cyclic mechanical strain (10% deformation, 0.15 Hz). Representative images were shown alongside quantitative analysis of DNA damage, expressed as the average number of 53BP1 foci per nucleus and the percentage of foci-positive nuclei (*n*=3-10). VSL#3 treatment did not significantly alter the extent of DNA damage under “Irr. All” conditions, while a significant reduction in DNA damage was observed in the “Irr. Medium” group with VSL#3 pre-treatment. Yellow bar, 20 μm; White bar, 50 μm. \**p*<0.05; \*\**p*<0.01; \*\*\**p*<0.001; ns, not significant.

## DISCUSSION

In this study, we leveraged a microengineered human gut-on-a-chip model to demonstrate GI-ARS and assess the effect of probiotic treatment. We observed that the gamma irradiation induced a morphological loss of the epithelial 3D structure only when the irradiated culture medium was perfused during the culture. This phenotypic outcome was linked to elevated ROS generation in the epithelial cells following perfusion of irradiated medium. Co-culturing of probiotic VSL#3 suppressed intracellular ROS and protected cells from morphological disruption. Thus, our gut-on-a-chip study positions VSL#3, and potentially other live probiotic bacteria, as a promising candidate for a medical countermeasure, offering a probiotic-based intervention that targets alleviating ROS-mediated damage to intestinal tissues. Importantly, we uncoupled the direct effects of radiation on epithelial cells from those mediated by reactive species, here ROS, in the surrounding microenvironment (i.e., culture medium), thereby facilitating an insight into the cytoprotective action of probiotic microbes. This uncoupling ability highlights the critical advantages of *in vitro* microphysiological models in minimizing confounding factors and generating new insights. Lastly, this approach offers an ethically preferable and practical alternative to nonhuman primate studies and human trials for radiation MCM.

High-energy photons, such as gamma rays, can ionize water molecules, a process called radiolysis, splitting water into highly reactive species like hydroxyl radicals (•OH), hydrogen radicals (H•), and hydrated electrons^55^. Thus, it can generate both extracellular ROS in the culture medium as well as intracellular ROS as cells are ∼70% water. Excessive ROS induces lipid peroxidation, protein oxidation, mitochondrial dysfunction, and DNA damage, which in turn lead to cell damage and death^56,57^. Although past MCM studies using organ-on-a-chip models have recreated and validated the radiation-induced ROS elevation *in vitro*^50,58^, the individual impacts of epithelial and medium irradiation have not been fully delineated.

We found that direct irradiation of epithelial cells induces DNA damage, cell injury, barrier impairment, and intracellular ROS. Furthermore, it can be deduced that the nuclear accumulation of 53BP1 foci in the “Irradiated All” group was mostly due to direct epithelial irradiation. The “Irradiated All” group also showed approximately 3.5 times stronger ROS signal compared to the “Irradiated Medium” group, implicating that direct epithelial irradiation contributed to most of the generated intracellular ROS. Thus, the presence of DNA damage and intracellular ROS generation could explain the elevated LDH levels in the “Irr. Cells” group. However, under a non-irradiated fresh medium perfusion, these epithelial damages through direct irradiation to the cells did not culminate in morphological disruption. A possible explanation is that a component of the culture medium, fetal bovine serum (FBS), contains albumin that has antioxidant properties by metal ligand binding and free radical scavenging^59^. Additionally, under the fresh medium perfusion, cell turnover could aid in shedding heavily damaged cells and removal through continuous medium flow. In contrast, continuous perfusion of ROS-rich irradiated medium in the “Irradiated Medium” and the “Irradiated All” group created a consistently hostile surrounding environment, causing significant morphological disruption of epithelial cells (regardless of direct irradiation to the cells), cell damage, and intracellular ROS, with nuclear DNA damage strongly elevated if cells were directly irradiated. Given the correlation between ROS generation and epithelial damage, it is crucial to understand whether ROS scavenging could protect the intestinal epithelial cells from radiation injury.

The VSL#3 is a probiotic formulation that includes 8 different bacterial strains (*Lactobacillus acidophilus*, *Lactiplantibacillus plantarum*, *Lacticaseibacillus paracasei*, *Lactobacillus delbrueckii* subsp. *bulgaricus*, *Bifidobacterium breve*, *B. longum*, *B. infantis*, *Streptococcus thermophilus*), where its anti-inflammatory and cytoprotective effects have been demonstrated in trials of various diseases^60^. In this study, VSL#3 administration, whether before or after radiation exposure, significantly preserved the epithelial 3D microarchitecture. This morphological protection of VSL#3 correlated with its ability to prevent intracellular ROS generation induced by combined epithelial and medium irradiation. The protective effect also corresponded with reduced LDH levels. We demonstrated that VSL#3 can effectively scavenge H_2_O_2_ at concentrations as low as 0.625 µM (Figure S4A). The reduction in ROS observed under both Irr. All and Irr. Medium conditions is likely attributable to the intrinsic antioxidant capacity of VSL#3 strains, potentially mediated by enzymatic mechanisms such as superoxide dismutase (SOD)^61^, catalase^62^, and peroxidase activity^63^. Given that the radiochemical yield (G-value) of H_2_O_2_ in pure water is 0.7-1.0 molecules/100 eV (0.0725–0.1036 µM/Gy)^64^, an 8 Gy dose would be expected to generate up to ∼0.8 µM H_2_O_2_ in the irradiated medium, assuming the culture medium is predominantly water. Based on our control experiment (Figure S4A), VSL#3 cells are capable of significantly reducing this ROS burden, likely through their demonstrated peroxidase activity (Figure S4B), thereby mitigating irradiation-induced oxidative stress. However, VSL#3 was unable to prevent double-strand breaks, likely due to the high penetrative power of gamma radiation. Despite the absence of ROS, radiation can directly break the chemical bonds in DNA^40^, evident by the almost 100% foci^+^ nuclei proportion of “Irradiated All”. In the “Irradiated Medium” group, we observed nuclear localization of 53BP1 foci only in the outer cells of the epithelial 3D microstructure, whereas inner cells did not have foci accumulation. We reason that peripheral cells, being the first exposed to extracellular ROS, developed both intracellular ROS and foci accumulation. When VSL#3 suppressed extracellular and intracellular ROS in both “Irradiated All” and “Irradiated Medium”, it enabled natural turnover of cells to aid in the shedding of damaged cells and maintenance of the 3D morphology. Although additional studies are needed to fully characterize this observation, it appears that the protective mechanism of VSL#3 is centered on mitigating oxidative stress rather than directly influencing the initiation of DNA damage.

Our findings align with previous gut-on-a-chip studies demonstrating the ability of VSL#3 to remarkably reduce oxidative stress^65^ and promote epithelial barrier function^51,66^ in inflammation and leaky gut models. *In vivo,* VSL#3 has been shown to attenuate ROS production by macrophages and reduce colonic injury in a murine colitis model^27^, as well as decrease ROS levels while increasing the antioxidant SOD in the brain tissue of a postoperative cognitive dysfunction mouse model^67^. Beyond VSL#3, a variety of probiotic candidates exhibit potent antioxidant and health-promoting properties. Traditional probiotics, such as *Lacticaseibacillus* and *Lactiplantibacillus* species, can mitigate oxidative stress^68^ through the production of SOD^69^ and catalase^70^. The microbial consortium of *Saccharomyces cerevisiae* and *Ketogulonigenium vulgare* produces 2-keto-L-gulonic acid (2-KLG), a precursor to vitamin C, which is a well-known antioxidant^71^, which in turn upregulates SOD and catalase in cultures. Engineering traditional probiotics to enhance the production of SOD and catalases^72^, or to biosynthesize antioxidant compounds such as lycopene^73^, represents a promising strategy for targeted functional enhancement. In addition, synbiotic formulations pairing a defined plant-based diet with urolithin-producing taxa such as *Gordonibacter*^74^ and *Enterocloster*^75^ could promote the generation of polyphenolic metabolites with broad range of antioxidant and anti-inflammatory activities^76,77^. Urolithins have been shown to reduce chemically induced colitis in mice^78^ and to mitigate radiation-induced intestinal injury^79^. Collectively, the exploration of traditional probiotics, engineered probiotics, and synbiotic strategies offers a comprehensive framework for developing live biotherapeutic countermeasures against radiation-induced damage through modulating oxidative stress.

Given the severe clinical consequences of radiation exposure^80,81^, the expansion of nuclear energy, and renewed commitments to space exploration^82^, there is an urgent global need for enhanced radiation preparedness. Governments are actively planning additional nuclear facilities to power artificial intelligence (AI) and other data-intensive technologies (Executive Order 14299), while the risk of nuclear incidents continues to pose a threat to civilian populations^83^. In space missions, protecting astronauts from cosmic radiation remains a major challenge^84^, as physical shielding alone is insufficient against high-energy cosmic rays, which can penetrate spacecraft and generate harmful secondary particles upon interactions with shielding materials^85^. These realities highlight the necessity not only of preventing radiation exposure but also of mitigating its biological impact through more effective medical interventions. The limited availability of FDA-approved radiomitigators hampers the ability to address the full spectrum of radiation-induced injuries, particularly in highly sensitive organs such as the brain and GI tract. Moreover, conventional animal models inadequately replicate human radiation responses^86^, and are increasingly constrained by ethical, regulatory, and logistical considerations^39^. These challenges are compounded by the high costs and operational barriers of conducting animal-based research in space environments^87^. Hence, these factors underscore the urgent need for physiologically relevant, human-specific platforms that are scalable, cost-effective, and experimentally controllable to advance the development of next-generation MCMs against radiation injury.

Organ-on-a-chip platforms have emerged as physiologically relevant *in vitro* systems that bridge the gap between basic research and translational medicine by enabling therapeutic evaluation in a human-relevant context. These microengineered systems enable controlled, reproducible studies that accelerate therapeutic screening and mechanism discovery under both terrestrial and spaceflight-relevant conditions^88,89^. For instance, gut-on-a-chip technology has been employed to demonstrate the protective effects of dimethyloxalylglycine, a prolyl hydroxylase inhibitor, as a potential mitigator of intestinal radiation injury^50^. Similarly, a lung alveolus-on-a-chip model identified heme oxygenase-1 as a promising target for mitigating pulmonary radiation damage^58^, while a bone marrow-on-a-chip model has elucidated the pathophysiological consequences of radiation on hematopoietic tissues^90^. Organ-on-a-chip technology has also advanced space biology research; for instance, a multi-organ microphysiological system linking bone marrow, cardiac, and hepatic tissues via a shared vascular circuit revealed that chronic neutron radiation induces apoptosis, stress responses, and cellular migration, while suppressing immune signaling in bone marrow-derived cells^91^. In this context, our finding that irradiated culture medium alone can cause significant structural and functional damage to the intestinal epithelium raises broader concerns about environmental exposure to radiation-contaminated resources, such as food and water. This observation highlights the urgent need for countermeasures that can mitigate indirect radiation effects, particularly in scenarios involving environmental contamination.

Taken together, our findings highlight the potential of organ-on-a-chip platforms as powerful preclinical testbeds for identifying and validating novel medical countermeasures against radiation-induced injuries. These human-relevant systems offer a unique opportunity to model complex physiological responses and hold significant promise for diverse applications, including protecting military personnel in combat environments, astronauts exposed to chronic cosmic radiation during space missions, and civilian populations during radiological or nuclear emergencies. In this study, we demonstrate that a commercially available probiotic mixture can mitigate radiation-induced intestinal injury by suppressing extracellular ROS, thereby preserving epithelial structure and function. This ROS-targeted mechanism suggests a promising breakthrough for probiotics as prophylactic interventions. Continued advancement and optimization of organ-on-a-chip technologies, particularly those incorporating human-derived samples and multi-organ integration, will be critical to enhance predictive validity and translational impact. These insights lay a foundation for future efforts to develop effective, accessible MCMs that address both direct and environmental radiation threats.

### Limitations of the study

Several limitations of our study warrant consideration and provide direction for future investigations. First, while the Caco-2 cell line used in this study originates from a human colorectal carcinoma, previous reports have shown that under physiologically relevant conditions within a gut-on-a-chip system, Caco-2 cells exhibit a heterogenous lineage and a transcriptome profile that is less cancer-like and more reflective of normal intestinal physiology^47,92^. Nonetheless, the use of human intestinal organoids derived from patient samples may offer enhanced fidelity by incorporating individual-specific genetic, epigenetic, and phenotypic characteristics^93^. Second, the short-term nature of our *in vitro* model that spans for several days post-radiation exposure limits its applicability to acute radiation injury, making it less suitable for modeling chronic radiation effects or long-term recovery dynamics. Third, our current model lacks a resident gut microbiota, which may influence the interaction and therapeutic outcomes of probiotics such as VSL#3. Incorporating a complex, pre-colonized microbial community would provide a more critically relevant context for evaluating microbiome-based interventions^94^. Finally, the absence of immune components restricts our ability to capture critical aspects of host-microbiome crosstalk, which are likely to influence the pathophysiology of GI ARS and the efficacy of live biotherapeutic products. Future studies integrating immune cells and other supporting cell types will be essential to more accurately recapitulate the intestinal microenvironment and to refine the predictive power of gut-on-a-chip platforms for translational applications.

## RESOURCE AVAILABILITY

### Lead Contact

Further information and requests for resources and reagents should be directed to and will be fulfilled by the Lead Contact, Hyun Jung Kim.

### Materials availability

Materials generated in this study are available from the Lead contact with a completed Materials Transfer Agreement.

### Data and code availability

Data available upon reasonable request to the corresponding author.

## ACKNOWLEDGEMENTS

This work was supported in part by the Food & Drug Administration (# 75F40119C10098 to D. E. I.) and the Kenneth Rainin Foundation Innovator Award (to H.J.K.). We thank Elisabeth Jiang and Amanda Jiang (Wyss Institute, Harvard) for their assistance in irradiation experiment. Figures were partially created in BioRender.com.

## AUTHOR CONTRIBUTIONS

D.E.I. and H.J.K. designed the study. H.J.K. and T.M. performed experiments. N.T. and H.J.K. analyzed the data and wrote the manuscript. All authors reviewed and approved the final version of the paper.

## STAR METHODS

Detailed methods are provided in the online version of this paper and include the following:

- KEY RESOURCES TABLE
- EXPERIMENTAL MODEL AND STUDY PARTICIPANT DETAILS

- Microfluidic human gut-on-a-chip model
- METHOD DETAILS

- Microfabrication of gut-on-a-chip device
- Gamma irradiation
- Host-probiotic co-cultures
- Assessment of DNA damage
- Evaluation of epithelial cell death
- Evaluation of epithelial barrier function
- ROS quantification
- Evaluation of H2O2 scavenging activity
- Microscopic analysis
- QUANTIFICATION AND STATISTICAL ANALYSIS

Document S1. Figures S1–S5

## START METHODS

KEY RESOURCES TABLE

| REAGENT or RESOURCE | SOURCE | IDENTIFIER |
| --- | --- | --- |
| Antibodies |  |  |
| Rabbit Polyclonal 53BP1 antibody | Abcam | ab36823 (RRID: AB_722497) |
| Goat Anti-Rabbit IgG H&L (DyLight® 488) | Abcam | ab96883 (RRID: AB_10679533) |
| Bacterial strains |  |  |
| VSL#3 (contains *Lactobacillus acidophilus*, *Lactiplantibacillus plantarum*, *Lacticaseibacillus paracasei*, *Lactobacillus delbrueckii* subsp. *bulgaricus*, *Bifidobacterium breve*, *Bifidobacterium longum*, *Bifidobacterium infantis*, *Streptococcus thermophilus*) | VSL Pharmaceuticals Inc. | VSL#3 Powder |
| Chemicals, peptides, and recombinant proteins |  |  |
| SYLGARD 184 Silicone Elastomer Kit | Dow Corning | 4019862 |
| Rat tail collagen type I | Gibco | A10483-01 |
| Matrigel | Corning | 356235 |
| Trypsin/EDTA solution (TE) | Gibco | R001100 |
| L-cysteine | Sigma Aldrich | C1276 |
| Bovine serum albumin | Sigma Aldrich | A9418 |
| Paraformaldehyde, 4% (w/v) | Electron Microscopy Science | 157-4 |
| Triton X-100 | Sigma Aldrich | 97062-208 |
| 4',6-diamidino-2-phenylindole dihydrochloride (1 mg/mL) | Thermo Scientific | 62248 |
| Critical commercial assays |  |  |
| CytoTox 96 Non-Radioactive Cytotoxicity Assay | Promega | G1780 |
| H2DCFDA, SE | Invitrogen | D2935 |
| CellROX Orange | Invitrogen | C10443 |
| Amplex Red Hydrogen Peroxide/Peroxidase Assay | ThermoFisher Scientific | A22188 |
| Experimental models: Cell lines |  |  |
| Human: Caco-2 BBE | Harvard Digestive Disease Center | Caco-2 BBE (RRID: CVCL_1096) |
| Software and algorithms |  |  |
| ImageJ 1.53t | National Institute of Health | https://imagej.nih.gov/ij/ |
| GraphPad Prism 10.1.2 | GraphPad Software | https://www.graphpad.com/ |
| LAS X | Leica Microsystems | https://www.leica-microsystems.com/products/microscope-software/p/leica-las-x-ls/ |

## EXPERIMENTAL MODEL AND STUDY PARTICIPANT DETAILS

### Microfluidic human gut-on-a-chip model

Human intestinal epithelial Caco-2 BBE cells (Harvard Digestive Disease Center) were cultured in T75 flasks using Dulbecco’s Modified Eagle Medium (DMEM; Gibco) supplemented with 20% (v/v) fetal bovine serum (FBS; Gibco), 100 U/mL penicillin and 100 μg/mL streptomycin (both from Gibco). Upon reaching 90% confluency, cells were detached using a 0.25% (v/v) trypsin and 0.02% (w/v) EDTA solution (Gibco), and subsequently resuspended in a complete medium at 5×10^6^ cells/mL. Prior to cell seeding, the microchannels of a gut-on-a-chip device were activated via UV/ozone treatment for 1 h and coated with an ECM mixture of 1% (v/v) Matrigel (Corning) and 30 μg/mL collagen type I (Gibco). The prepared cell suspension was introduced into the apical microchannel, and cells were allowed to adhere for 1 h under static condition in a humidified incubator (37°C, 5% CO_2_). Following initial attachment, perfusion was initiated through the apical channel at 30 μL/h for 24 using a syringe pump (Braintree Scientific). Thereafter, bidirectional flow was established in both apical and basolateral channels at 30 μL/h, producing ∼0.02 dyne/cm^2^ shear stress. Once epithelial confluence and tight junction integrity were confirmed, cyclic mechanical strain (10% cell strain at 0.15 Hz frequency) was applied using a computer-controlled vacuum system (Flexcell International Corporation) connected to the side vacuum chambers. These physiodynamic conditions of flow and deformations were maintained throughout irradiation exposures and co-culture experiments.

## METHOD DETAILS

### Microfabrication of a gut-on-a-chip device

The gut-on-a-chip device was fabricated using standard soft lithography techniques, as previously described^46,49^. Silicon molds for the upper and lower microchannels (dimensions: 150 μm height × 1000 μm width × 10 mm length) were fabricated via photolithography.

Polydimethylsiloxane (PDMS; Sylgard 184, Dow Corning) was prepared by mixing the elastomer base and curing agent at a 15:1 (w/w) ratio, followed by degassing under vacuum. The mixture was poured onto the silicon molds and cured at 60°C for at least 4 h. After curing, the PDMS layers were demolded, and inlet/outlet ports (2 mm diameter) were punched using a Harris Uni-Core punch (GE Healthcare). The porous PDMS membrane was fabricated using a micropatterned silicon wafer featuring cylindrical micropillars (10 μm diameter, 25 μm center-to-center spacing, 25 μm height). Approximately 10 g of degassed PDMS (15:1 ratio, w/w) was poured over the patterned wafer and covered with a pre-silanized PDMS slab with a weight (∼3 kg) to ensure uniform thickness. The setup was cured overnight at 60°C to form a thin, porous membrane. Device assembly was performed in sequential bonding steps. First, the upper PDMS microchannel layer and the porous membrane were exposed to oxygen plasma (COVANCE, Femto Science Inc.; 125 W and 20 Pa for 75 seconds) and brought into conformal contact to facilitate irreversible bonding, followed by baking at 80°C for 4 h. The bonded upper layer-membrane assembly was then aligned under a stereoscope (MDG41, Leica Microsystems) with the plasma-activated lower PDMS channel layer and sealed to form the complete device. Fluidic access was established using 18-gauge blunt-end stainless steel needles (Shintop) bent at 90°, trimmed to the appropriate length, and inserted into the pre-punched ports. Each needle was connected to a 2-inch segment of silicone tubing (Tygon 3350, 1/32“ ID, 3/32” OD, Saint-Gobain) to enable integration with syringe pumps and peripheral flow control system.

### Gamma irradiation

Gut-on-a-chip devices containing 3D undulated intestinal epithelial layers were disconnected from the syringe pump and promptly transported to the irradiation facility. Devices were subjected to a single 8 Gy dose of gamma (γ) irradiation using a cesium-137 (Cs-137) source (Gammacell 40 Exactor) at a dose rate of 0.78 Gy/min for 10.3 min. Throughout the irradiation procedure, the chamber temperature was maintained at 37°C to preserve physiological conditions. Control devices underwent the same handling and transportation procedures but were not exposed to irradiation. Separately, culture medium alone was irradiated under identical conditions to generate the “Irradiated Medium” group for downstream experiments.

### Host-probiotic co-cultures

To perform host-probiotic co-cultures, VSL#3 powder (VSL Pharmaceuticals Inc.) was inoculated in 3 mL of a 1:1 (v/v) mixture of pre-autoclaved Lactobacilli MRS Broth (MRS; Difco) and Reinforced Clostridial Medium (RCM; Difco) in a 15 mL test tube. The bacterial culture was incubated anaerobically for 12-18 h without shaking using a GasPak EZ container (BD Diagnostics) equipped with GasPak EZ sachets. Following enrichment, the bacterial suspension was centrifuged (10,000×*g*, 5 min), and the pellet was resuspended in an antibiotic-free cell culture medium supplemented with 1 mg/mL L-cysteine (Sigma Aldrich) to promote an oxygen-reducing environment. The final density of viable probiotic cells was adjusted to 1×10^7^ CFU/mL. Prior to bacterial inoculation, gut-on-a-chip devices were pre-conditioned with an antibiotic-free medium containing L-cysteine (1 mg/mL) for at least 12 h. The prepared probiotic suspension was introduced into the apical channel of the device, allowing bacterial adhesion to the apical epithelial surface during a 1-hour static incubation at 37°C and 5% CO_2_. After adhesion, Physiomimetic conditions were restored by initiating continuous flow (40 μL/h) and cyclic mechanical strain (10% strain, 0.15 Hz) for the duration of the experiment. Probiotic co-culture was conducted either 24 h prior to irradiation (Pre-VSL#3) or immediately following irradiation (Post-VSL#3), and maintained for a minimum of 48 h to evaluate host-microbe interactions under physiologically relevant conditions.

### Assessment of DNA damage

DNA double-strand breaks were visually assessed via immunofluorescence staining of 53BP1, a nuclear marker of DNA damage. Cells in the gut-on-a-chip device were fixed with 4% (w/v) paraformaldehyde (PFA; Electron Microscopy Science) for 20 min, permeabilized with 0.3% (v/v) Triton X-100 (Sigma Aldrich) for 20 minutes, and blocked with filter-sterilized 2% (w/v) bovine serum albumin (BSA; Sigma Aldrich) for 1 h at room temperature. Cells were rinsed with PBS between steps. Rabbit anti-53BP1 primary antibody (Abcam), resuspended in 2% BSA, was introduced to both microchannels and incubated statically for 1 h at room temperature. After PBS washing, DyLight 488-conjugated goat anti-rabbit secondary antibody (Abcam), prepared in 2% BSA, was added and incubated statically for 1 h at room temperature under light protection. Nuclei were counterstained with 4’,6-diamidino-2-phenylindole (DAPI, 5 µg/mL; Molecular Probe) for 30 minutes. Following final PBS washes, samples were imaged using confocal laser scanning microscope (SP5 X MP DMI-6000; Leica). DNA damage was quantified by two metrics: 1) the average number of 53BP1 foci per nucleus, and 2) the percentage of nuclei containing one or more 53BP1 foci relative to total number of DAPI-positive nuclei. Only foci localized within nuclear boundaries were included in the analysis.

### Evaluation of epithelial cell death

Epithelial cell death was assessed by measuring lactate dehydrogenase (LDH) activity using the CytoTox 96 Non-Radioactive Cytotoxicity Assay kit (Promega), following the manufacturer’s protocol. Briefly, effluents from both apical and basolateral microchannels were collected at designated time points before and after irradiation. Samples were centrifuged at 10,000×*g* for 5 min to remove debris, and the supernatants were stored at -80°C until analysis. For the assay, thawed samples were incubated with LDH substrate reagent for 30 min at room temperature, followed by the addition of a stop solution to terminate the reaction. Absorbance was measured at 492 nm using a microplate reader (SpectraMax M5; Molecular Devices). To determine the maximum LDH release, control cells were lysed using lysis solution provided in the kit prior to substrate addition. Percent cytotoxicity was calculated by normalizing background-corrected sample absorbance values to the maximum LDH release control. This quantitative measurement provided an estimate of cell membrane integrity and death in response to experimental conditions.

### Evaluation of epithelial barrier function

Epithelial barrier integrity was assessed by measuring transepithelial electrical resistance (TEER) across the cell layer within the gut-on-a-chip device. Two Ag/AgCl electrodes (0.008′′ in diameter; A-M systems, Inc.) were inserted into the upper and lower microchannels, and resistance was recorded using a digital multimeter (87V Industrial Multimeter; Fluke Corporation). A cell-free gut-on-a-chip device filled with culture medium served as the baseline control to account for background resistance. TEER values (kΩ cm^2^) were calculated using the formula: TEER = (Ω_t_ − Ω_blank_) × A, where Ω_t_ is the measured resistance at the experimental timepoint, Ω_blank_ is the resistance of cell-free control chip, and A is the surface area of the cell culture (∼0.11 cm^2^ in this system).

### ROS quantification

Extracellular ROS levels within the irradiated medium were quantified using the H2DCFDA fluorogenic probe (Invitrogen), following the manufacturer’s instructions. Fluorescence intensity was measured using a microplate reader (SpectraMax M5; Molecular Devices). Intracellular ROS production was assessed using CellROX Orange (Invitrogen), a cell-permeable fluorogenic dye that selectively detects cytoplasmic free radicals. The reagent was diluted 1:250 in culture medium and perfused through both microchannels at 30 μL/h (37°C, 5% CO_2_) for 1 h. After PBS washing, cells were imaged using confocal laser scanning microscopy (Leica SP5 X MP DMI-6000). Mean fluorescence intensity per field of view was quantified using ImageJ software.

### Evaluation of H_2_O_2_ scavenging activity

The ability of VSL#3 to scavenge H_2_O_2_ and exhibit peroxidase activity was assessed using the Amplex Red Hydrogen Peroxide/Peroxidase Assay Kit (A22188; Thermo Fisher Scientific). Briefly, VSL#3 cells were anaerobically pre-enriched in antibiotic-free DMEM supplemented with 20% (v/v) FBS overnight without shaking at 37 °C. The cultured microbial cells were washed twice by repeating centrifugation (5000×*g* for 5 minutes) and washing steps with the same fresh medium, then resuspended in the medium to a final optical density at 600 nm of approximately 0.8. For the H_2_O_2_ scavenging assay, serially diluted H_2_O_2_ standard solutions (0-20 µM) served as controls. A separate series containing H_2_O_2_ (0-40 µM) was mixed with VSL#3 suspension (1:1, v/v; final H_2_O_2_ concentration, 0-20 µM). Each mixture was then combined 1:1 (v/v) with 100 µM Amplex Red reagent and 0.2 U/mL horseradish peroxidase (HRP) and incubated at 37 °C for 30 min. For the peroxidase activity assay, serially diluted HRP standards (0-2 mU/mL) or VSL#3 suspension alone were combined 1:1 (v/v) with 100 µM Amplex Red reagent and 2 mM H_2_O_2_, followed by incubation at 37 °C for 30 min. Fluorescence was measured using a plate reader (Cytation 5, BioTek) at 545 nm excitation and 590 nm emission.

### Microscopic analysis

Three-dimensional epithelial structures were imaged using both phase contrast and differential interference contrast (DIC) microscopy. Phase contrast micrographs were captured using Axiovert 40CFL microscope (Zeiss) equipped with a 20× Ph1 objective (NA 0.30; Zeiss) and a Moticam 2500 camera (Motic China Group Co., Ltd.), operated with Motic Images Plus 2.0 software. For higher-resolution imaging, DIC microscopy was performed using an Axio Observer.Z1 microscope (Zeiss) coupled to a 20× LD PlnN DICII objective (NA 0.4; Zeiss) and a high-resolution digital camera EM-CCD digital camera (C9100; Hamamatsu, 1,000 × 1,000 pixels, 8 µm pixel size) using either MetaMorph (Molecular Devices) or ZEN Pro (Zeiss) software for image acquisition. Epithelial morphology, including villi-like structures, was assessed by imaging more than 10 distinct regions per sample from at least two independent gut-on-a-chip replicates at each time point. Representative images were selected for figure presentation. For confocal imaging, samples were analyzed using a laser scanning confocal microscope (Leica SP5 X MP DMI-6000). The 3D epithelial microarchitecture was visualized via immunofluorescence using a 25× water-immersion objective (NA 0.95; Leica) linked to a 405-nm diode laser and a tunable white light laser (489-670 nm), with detection through photomultiplier tube or HyD detectors. Single-plane fluorescence images were acquired and processed using Leica LAS AF software (Ver. 2.3.1, build 5194; Leica Microsystems).

## QUANTIFICATION AND STATISTICAL ANALYSIS

All data and error bars in the article are represented as mean ± standard error of the mean (SEM). Replication number, *n*, is indicated in each figure legend. We applied a two-tailed unpaired t-test (Figures 1D and 3A-C), multiple unpaired t-test followed by a false discovery rate (FDR) correction (Figure 2C), an ordinary one-way analysis of variance (ANOVA) followed by a Dunnett correction (Figures 3B-C, 4B, and S4B), or a two-way ANOVA followed by a Sidak correction (Figures 1C, 2B, 4C-D, 5, S3, and S4A). Differences between groups were considered statistically significant when *p*, corrected *p*, or FDR < 0.05. All statistical analyses were carried out using GraphPad Prism 10.1.2 (GraphPad Software).

