## Supplemental document 1 for "Probiotic Intervention Mitigates Radiation-Induced Intestinal Injury by Alleviating Oxidative Stress in a Human Gut-on-a-chip"

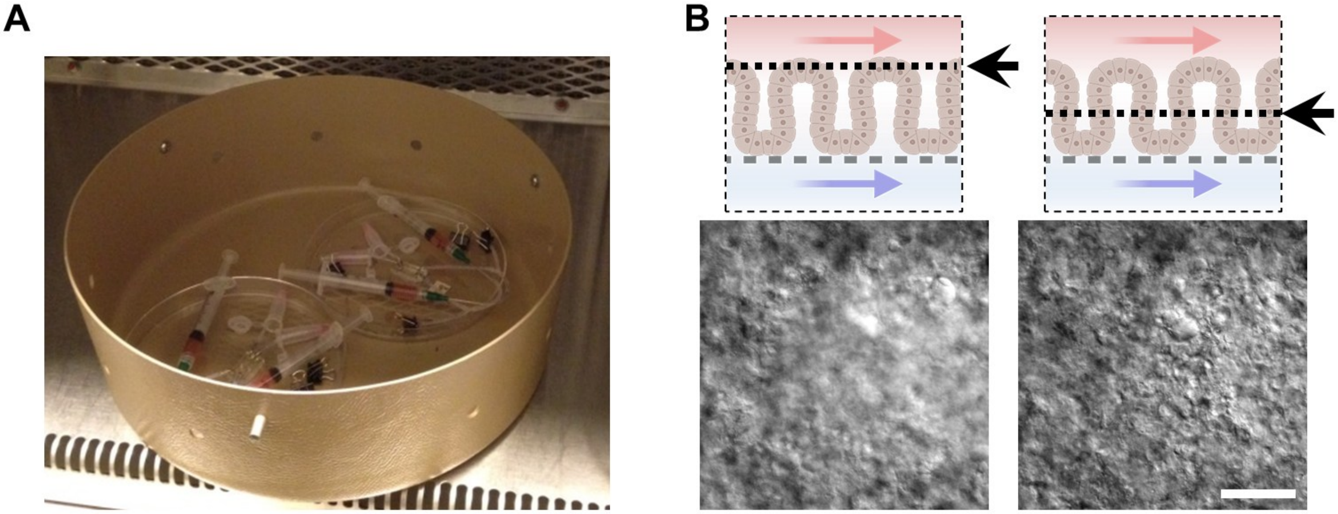

**Figure S1. Gamma irradiation setup and post-irradiation morphological assessment.** (A) Representative gamma irradiation setup showing two gut-on-a-chip devices placed within the round chamber of the Gammacell 40 Exactor. For “Irradiated Medium” and “Irradiated All” experiments, culture medium in a 50-mL tube was irradiated under identical conditions. For “Irradiated Cells” experiments, irradiated syringes were replaced with non-irradiated ones. (B) Schematic illustration indicating the focal plane position and corresponding representative DIC micrographs of post-irradiation morphologies. The grey dashed line in the schematic denotes the basement membrane location. Arrows indicate the direction of culture medium flow in the upper and lower microchannels. Bar, 100 µm.

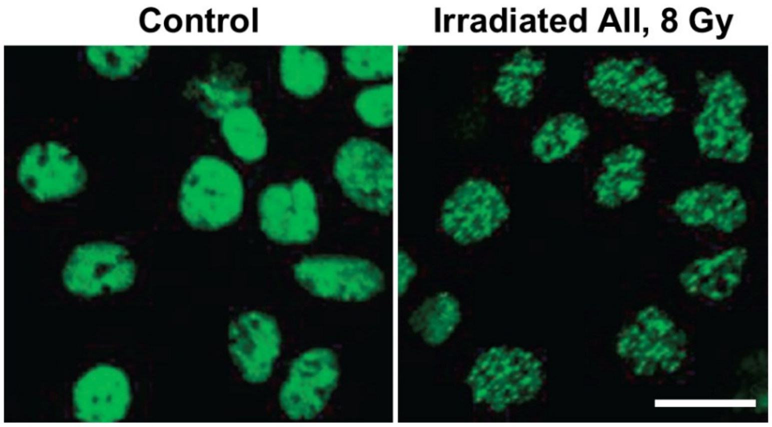


**Figure S2. Formation of double-strand DNA damage foci in a 2D Caco-2 monolayer following irradiation under static culture conditions.**

A Caco-2 monolayer pre-cultured on an ECM-coated glass-bottom plate under static conditions was exposed to 8 Gy irradiation, followed by incubation at 37 $^{o}$C in a $CO_2$ incubator for 5 h with irradiated medium. Cells were then immunofluorescently labeled with anti-53BP1 antibodies. The presence of multiple foci indicates sites of double-strand DNA breaks. Bar, 20 μm.

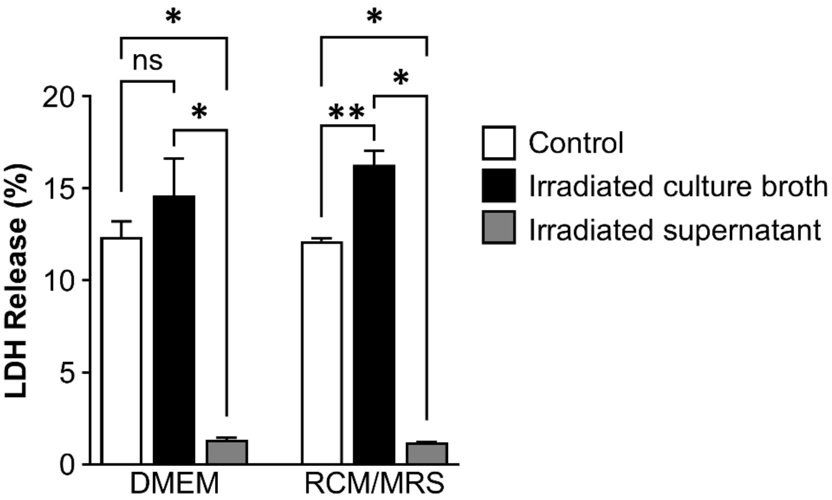


**Figure S3. Background LDH signals detected in probiotic VSL#3 cultures.**

Culture broths of VSL#3 cells grown in either DMEM or RCM/MRS bacterial medium were assessed to determine background LDH activity. “Control” represents the culture broth of VSL#3 cells (final density, $1\times10^{7}$ CFU/mL) grown in each medium. “Irradiated culture broth” denotes the samples of the whole VSL#3 culture broth irradiated at 8 Gy in each medium. “Irradiated supernatant” refers to the irradiated samples (8 Gy) prepared from culture supernatants obtained by centrifugation (10,000× *g*, 5 min) of each culture broth. Data are shown as mean ± SEM ($n$=3). *$p$<0.0001; **$p$<0.05.

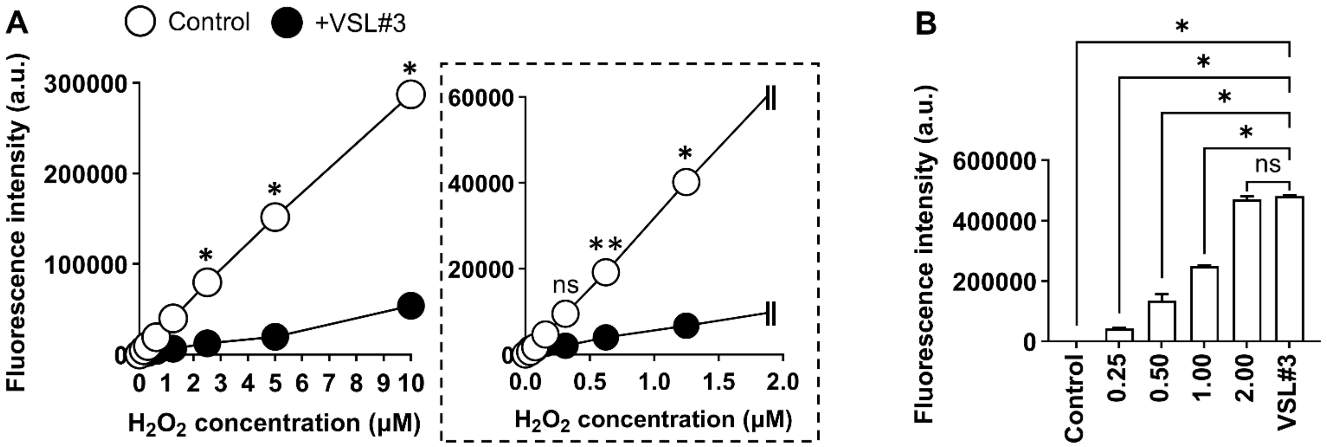


**Figure S4. Probiotic VSL#3 exhibited $H_2O_2$ scavenging and peroxidase activities.**

(A) Fluorescence quantification of reactions between 0.2 U/mL horseradish peroxidase (HRP) and 100 µM Amplex Red reagent with varying $H_2O_2$ concentrations in the absence or presence of VSL#3 cells (*n* = 2). The dashed inset highlights the lower concentration range; no statistically significant differences were observed for concentrations <0.5 µM. Fluorescence intensity is directly proportional to $H_2O_2$ concentration. $^*p<0.001$; $^{**}p<0.05$; ns, not significant. (B) Fluorescence quantification of reactions between 2 mM $H_2O_2$ and 100 µM Amplex Red reacting with either varying HRP concentrations (0-2.00 mU/mL) or VSL#3 (*n* = 2). Fluorescence intensity is directly proportional to peroxidase concentration. $^*p<0.001$; ns, not significant. The Control reaction contains neither HRP nor bacteria.

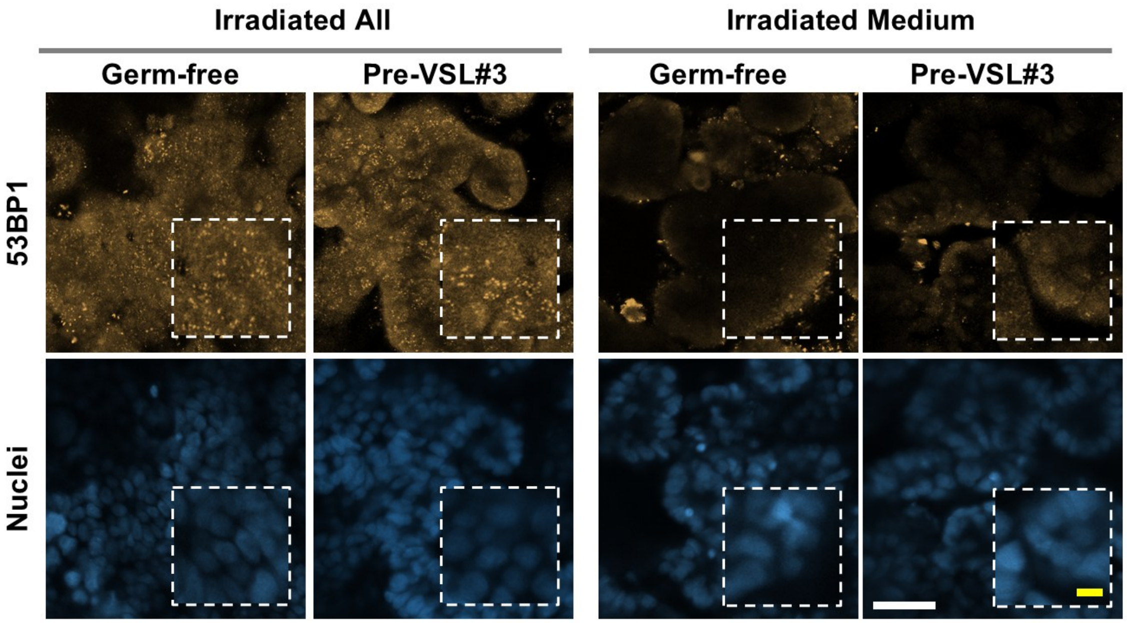


**Figure S5. Probiotic VSL#3 does not alter radiation-induced DNA damage response.** Representative immunofluorescence micrographs showing independent signals of 53BP1 and Nuclei in 3D Caco-2 cells subjected to either Irradiated All (direct irradiation followed by exposure to irradiated medium) or Irradiated Medium (intact Caco-2 cells exposed to irradiated medium), in the presence or absence of VSL#3 cells. The individual fluorescence channels corresponding to the overlaid images in Figure 5 are shown separately. Yellow bar, 20 μm; White bar, 50 μm.
